# eQTL mapping in transgenic alpha-synuclein carrying *Caenorhabditis elegans* recombinant inbred line

**DOI:** 10.1101/2023.08.18.553811

**Authors:** Yuqing Huang, Yiru A. Wang, Lisa van Sluijs, Demi H. J. Vogels, Yuzhi Chen, Vivian I. P. Tegelbeckers, Steven Schoonderwoerd, Joost A.G. Riksen, Jan E. Kammenga, Simon C. Harvey, Mark G. Sterken

**Affiliations:** Laboratory of Nematology, Wageningen University & Research, Wageningen, the Netherlands; Faculty of Engineering and Science, University of Greenwich, United Kingdom

**Keywords:** α-synuclein, *C. elegans*, RIL, IL, eQTL, genetic variation

## Abstract

Protein aggregation of α-synuclein (αS) is a genetic and neuropathological hallmark of Parkinson’s disease (PD). Studies in the model nematode *Caenorhabditis elegans* suggested that variation of αS aggregation depends on the genetic background. However, which genes and genetic modifiers underlie individual differences in αS pathology remains unknown. To study the genotypic-phenotypic relationship of αS aggregation, we constructed a Recombinant Inbred Line (RIL) panel derived from a cross between genetically divergent strains *C. elegans* NL5901 and SCH4856, both harboring the human αS gene. As a first step to discover genetic modifiers 70 αS-RILs were measured for whole-genome gene expression and expression quantitative locus analysis (eQTL) were mapped. We detected multiple eQTL hot-spots, many of which were located on Chromosome V. To confirm a causal locus, we developed Introgression Lines (ILs) that contain SCH4856 introgressions on Chromosome V in an NL5901 background. We detected 74 genes with an interactive effect between αS and the genetic background, including the human p38 MAPK homologue *pmk-1* that has previously been associated with PD. Together, we present a unique αS-RIL panel for defining effects of natural genetic variation on αS pathology, which contributes to finding genetic modifiers of PD.

## Introduction

Protein aggregation is a key process in the pathogenesis of a number of human diseases, including neurodegenerative diseases (NGDs) (1) such as Alzheimer Disease (AD) and Parkinson’s Disease (PD) (2). NGDs affect neurons of the central nervous system (CNS) and are associated with characteristic clinical symptoms including dementia and motor disability (3,4). NGDs have the pathological hallmark of chronic, progressive multi-system neurodegenerative effects that mainly affect the elderly (5,6). Due to the increasing aging of human populations worldwide, there is a pressing need for a deeper molecular and genetic understanding of NGDs where protein aggregation is one of the most prominent pathogenesis-related factors (7).

Protein aggregation has been found to impact organisms at various levels, from effects on cellular organization up to distinct anatomical features. For example, aggregated α-synuclein (αS) in PD and aggregated amyloid in AD may cause loss-of-function or toxic gain-of-function in the cell (8,9). These aggregations trigger tissue abnormalities related to disease pathology and play an important role in the molecular cascade of pathogenicity. At the cellular level, these protein aggregations interrupt proteostasis and may trigger toxicity pathways, which can be influenced by age and physiological conditions (10). At the tissue level, both extracellular and intracellular aggregates are involved in pathogenesis, including amyloid plaques and neurofibrillary tangles in the affected human brain (8). Genetic variation plays a critical role in the onset and progression of NDGs (11) and has been at the heart of quantitative genetic studies aiming to identify and characterize causal loci associated with pathologies of NDGs. However, the detection of genetic modifiers and their functional role in NDGs phenotypes has been hampered because of the poor genetic tractability of humans. This can be overcome by studying the genetic variation of model organisms in which genes can be easily knocked out or overexpressed in different genetic backgrounds (12,13). For instance, the polymorphic monoamine oxidase A coding gene *amx-2* was identified as a major modifier of the oncogenic RAS/MAPK signaling pathway (14). Subsequently, it was found that *amx-2* also was causal as gene expression regulator (eQTL) associated with Ras/MAPK signaling (15). These studies were conducted in the model nematode *C. elegans* for which powerful genetic tools are available for studying genetic variation (16–21).

In *C. elegans* there is ample natural genetic variation that is accessible to researchers because of the availability of genetically diverse strains (Cook *et al.*, 2017) and the level of genetic diversity of *C. elegans* is close to that of humans (22). In addition, in the genes of *C. elegans* that encode proteins, ∼38% was estimated to have human orthologs (23). *C. elegans* also has the advantages of a large number of progeny, a rapid life cycle and short lifespan, transparent body and the ease of genetic modification (24). Importantly, its advantages allow for the construction of recombinant inbred lines (RILs) derived from genetically diverse strains, to obtain a large number of progeny with different genetic backgrounds. *C. elegans* RILs have often been used in many quantitative trait loci (QTL) mapping studies applied to stress-response, lifespan analysis and metabolic studies(25–27). QTL mapping in RILs facilitates the study of how genetic variation influences αS aggregation across different genetic backgrounds.

Here, we generated the αS-RILs carrying an αS introgression by reciprocal crosses between NL5901 and SCH4856 (28) to identify genetic locations associated with genetic modifiers of αS related phenotypes. We present the genetic map of the panel and measure gene-expression and estimated the heritability and transgressive segregation of gene expression. Furthermore, we characterize the effect of local (*cis*) and distant (*trans*) genetic variation on gene expression changes. Using this approach we uncover the genetic architecture and main hotspots for genetic modifiers of αS phenotypes in an αS-RIL population followed by conformation with introgression lines (ILs.)

## Methods

### Nematode culturing and strains

*C. elegans* strains N2, CB4856, NL5901, SCH4856, and N2×CB4856 ILs WN268, WN269, and WN270 (14,29) were kept at 20°C on Nematode Growth Medium (NGM) plates seeded with *Escherichia coli* OP50 as food resource (30). To age-synchronize the nematodes, we transferred starved nematode populations to fresh NGM plates, allowed for egg laying (after three days), and then used bleach to dissolve the adults and keep the eggs. Next, the eggs were transferred to fresh 9cm NGM plates.

### Construction of the αS-RIL population and αS-ILs

The αS-RILs were constructed by crossing NL5901 and SCH4856 in two phases, ultimately leading to a panel of 212 RILs, of which 90 were genotyped by PCR, and 88 out of 90 were sequenced.

During the first phase, NL5901 hermaphrodites were crossed with SCH4856 males and NL5901 males crossed with SCH4856 hermaphrodites, to ensure F1 generation inherited mitochondria from both genetic backgrounds. Subsequently, the F1 generation was backcrossed with SCH4856 males to obtain a mix of alleles at the *peel-1/zeel-1* incompatibility locus on chromosome I (31). From these crosses, single hermaphrodites were selected and propagated and inbred further. After the next eight generations of inbreeding by self-fertilization, 93 RILs were obtained. However, partial genotyping by PCR and gel electrophoresis showed that these RILs displayed a low rate of crossovers and had a skewed N2/CB4856 distribution, probably because of poor male performance during mating.

To construct a more diverse RIL panel, we selected eight (out of 93) genetically diverse RILs and NL5901 and SCH4856, which were used for a new round of crossing. We first inter-bred the strains, generating a pool of 45 F1 populations, which were subsequently randomly crossed. Also the F2 and the F3 generations were crossed in a random manner, keeping the number to 45 heterozygous populations. After the F3 cross, the number of populations was expanded to approximately 200 populations (singled out hermaphrodites), which were propagated by self-fertilization for another eight generations by transferring single hermaphrodites to new plates. At the end, a population of 212 homozygous αS-RILs were obtained, including the eight founder strains.

#### αS-RIL genotyping and sequencing

Out of the 212 αS-RILs, 90 RILs were randomly selected for PCR-based genotyping to construct a rough genetic map. From each RIL, nematodes were collected for PCR-based genotyping with primers detecting insertions/deletions (Indels) between the N2 and CB4856 genomes (28). PCRs were conducted using the GoTaq^®^ DNA polymerase kit and 41 pairs of primers covering the arms and central area of each chromosome were used (**Supplementary Table S1**) (32). For this step, nematodes were lysed at 65°C for 30 minutes with a custom lysis buffer, followed by 5 minutes at 99°C. PCR amplification was performed with an annealing temperature of 58°C (30 seconds) and an extension time of 1 minute for 40 cycles. Amplicons were visualized on 1.5% agarose gels stained with ethidium bromide. This step was used as verification of the crosses.

After being assured a genetically divergent set of strains was created, 88 out of 90 RILs were picked together with parental and reference strains for whole-genome DNA sequencing to construct a high-resolution genetic map. Before DNA sequencing, DNA was extracted using the DNeasy® 96 Blood & Tissue Kit (QIAGEN, Germany). The DNA concentration was measured using Qubit 3.0 Fluorometer (Thermo Fisher Scientific). For each strain, a total of 0.75 ng of DNA was combined with 2.5 µl 1/35th diluted transposome (Illumina; purchased in kit # FC-121-1011), and 1X Tris Buffer (10X Tris Buffer: 100 mM Tris-HCl pH 8.0, 50 mM MgCl_2_) in a 10 µl final volume on ice. This reaction was incubated at 55°C for 10 minutes. The amplification reaction for each strain contained (final concentrations): 1X ExTaq Buffer, 0.2 mM dNTPs, 1 Unit ExTaq (Takara, catalog #RR001A), 0.2 µM primer 1, 0.2 µM primer 2, and 5 µl of material from the previous step in a 25 µl total volume. Each strain had a unique pair of indexed primers. Amplification was carried out in a thermocycler with the following conditions: 72°C for 3 minutes (1X); 95°C for 30 seconds (1X); 95°C 10 seconds, 62°C 30 seconds, 72°C 3 minutes (20X); 10°C hold. 16 µl from each amplification reaction was used to generate a pool of 48 libraries. We electrophoresed the libraries on a 2% agarose gel, excised DNA in the range of 500-700 bp, and gel purified the sample with Qiagen’s Gel Purification Kit (catalog #28706). The resulting pooled samples were sequenced using the Illumina MiSeq platform.

#### Construction of the αS-RIL genetic map

The genotypes from the whole-genome HiSeq data were called as described previously (33). The raw data was deposited at array express (E-MTAB-12575). In short, variants were called versus the N2 genome with the Andersen Lab’s nil-ril nextflow pipeline (https://github.com/AndersenLab/nil-ril-nf). A hidden-markov-model (HMM) was used to fill in missing genotypes. The resulting vcf was used to generate a genotype matrix.

We integrated the sequencing-based genotype with the PCR-based genotype. This was done manually. First, we checked the primer genotype based on the DNA sequencing result: if the DNA sequencing genotype result for adjacent locations was contrary to the PCR-based genotype, we corrected either the DNA-sequencing result (if only a small area was called) or the PCR-based genotype (if there was strong evidence from the DNA-sequencing). Only few corrections were required. Second, as not all strains were genotyped with two methods, the map also contained blanks. These were filled in based on adjacent markers: if their genotypes were the same, we filled the intermediate blank with the same genotype; if their genotypes were not the same, then we left the blank. These steps resulted in a high-resolution genetic map with 1,927 markers.

#### Genetic map analysis

Our estimated genetic map for these 70 αS-RILs employed the rigorous selection of 124 mishybridization markers to ensure uniform coverage of the *C. elegans* genome. All the *cis*-eQTL were sorted by physical position, we further estimated genetic distances in αS-RILs. The data set includes: a) the counts of crossover breakpoints observed between each pair of markers in the genetic map; b) the unassigned genotype at the certain marker was predicted by the genotype of its closest surrounding marker. Then, the statistical analyses of recombination frequencies were performed in R (version 3.5.3 x64). Notably, most of chromosome IV was assigned as predominantly N2 due to both parental strains carrying the αS introgression (Wang *et al.* 2019).

#### IL construction

Introgression lines (ILs) were constructed to confirm eQTL *trans-*bands on Chromosome V. Three IL_N2_ strains, WN268, WN269, and WN270, were selected as parental lines because they carry a CB4856 introgression (in an N2 background) on Chromosome V (29). Hermaphrodites of these three strains were crossed with NL5901 males separately to obtain four genotypes of homozygous ILs: i) no-CB4856 introgression and no αS::YFP introgression (WN613, WN617, and WN621), ii) with a CB4856 introgression and no αS::YFP introgression (WN614, WN618, and WN622), iii) no CB4856 introgression and with an αS::YFP introgression (WN615, WN619, and WN623), and iv) with a CB4856 introgression and with an αS::YFP introgression (WN616, WN620, and WN624).

After generation of the F1, hermaphrodites were singled out for self-fertilization for 6 generations. Presence of the correct genotypes was confirmed in the F2. The presence of the introgression after crossing was determined by gel-electrophoresis using insertion-deletion primers (28). The presence of the αS transgene was done by observing YFP signal of the progeny under a stereo microscope equipped for YFP detection.

### Preparation of αS expressing strains

To test the expression dynamics, a timeseries microarray experiment was done on NL5901 and SCH4856 over the course of 48 up till 216 hours post synchronization (a sample was collected each 24 hours). First, the nematodes were age-synchronized by bleaching an egg-laying population. After approximately 48h of post-synchronization, nematodes were collected by rinsing the NGM plates with M9 buffer and then transferred to fresh NGM plates containing FUdR (Floxuridine, 200µl of 50mg/ml) to prevent newly-produced eggs from hatching (34). As such, nematodes on the plates were from the same generation. Subsequently, 96h post-synchronization, nematodes were transferred to fresh FUdR plates again to avoid starving. Otherwise nematodes would stop growing, and starvation might interfere with the expression. Upon sampling, nematodes were collected in Eppendorf tubes which were then centrifugated to gather all nematodes in a pellet. Finally, the supernatant was removed and the nematodes were flash frozen in liquid nitrogen and stored at −80°C until RNA isolation.

Subsequently, hermaphrodite nematodes were collected at 120h after bleaching for RILs expression sample collection. As the time series experiment indicated that mature adult worms present relatively stable expression levels, 120h old nematodes of all strains (from synchronized embryo onwards) were used as they do not go through age-dependent changes in gene expression anymore. The same experimental procedure was followed as described for the time-series. At and age of 120h nematodes were collected in Eppendorf tubes using M9 buffer, which were then processed and ultimately flash-frozen.

For the IL gene-expression experiment, the same setup as for the RILs was applied, where samples were also collected at an age of 120h. The experiment was conducted in three time-separated biological replicates whereby each replicate contained the full set of 16 ILs.

### Gene expression measurements

RNA isolation for the time series experiment, RIL experiment was done using the Maxwell® 16 AS2000 instrument with the Maxwell® 16 LEV simply RNA tissue kit . The IL experiment RNA isolation was conducted using the Maxwell® 16 LEV plant RNA kit (Promega) (35). The frozen nematodes were lysed using a modified lysis protocol including a proteinase K digestion (36). After isolation, the concentration was measured using Nanodrop (Thermofisher). RNA samples were stored at −80°C until cDNA synthesis.

The microarray scanning and data extraction was conducted as described by (36). Following ‘Two-Color Microarray-Based Gene Expression Analysis’ (version 6.0) protocol, the *C. elegans* (V2) Gene Expression Microarray 4 × 44k chips were used (Agilent). Scanning was done with an Agilent High Resolution C Scanner. For extraction, we used Agilent Feature Extraction (version 12.1.1.1) based on the manufacturer’s guidelines. Raw data was deposited on ArrayExpress, under E-MTAB-11657 (timeseries experiment), E-MTAB-11658 (RIL experiment), and E-MTAB-11661 (IL experiment).

#### RT-qPCR for mRNA abundance of αS

RNA was extracted from the frozen nematode sample pellets. For lysis step, proteinase K was added for digestion (37), and then RNA isolation was done by Maxwell® AS2000 (Promega) using the Maxwell® 16 LEV plant RNA kit (Promega). The RNA concentration was measured by Nanodrop (Thermofisher). RNA samples were stored at −80°C or applied directly for cDNA synthesis.

RT-qPCR was started by cDNA synthesis. According to the manufacturer’s protocol (Promega), RNA was reverse transcribed to cDNA firstly with random hexamers (50 ng/µl) and dNTPs (5mM) and then by adding GoScript Reaction Buffer (5×), MgCl_2_ (25 mM), RNasin and GoScript Reverse Transcriptase. Samples were used for qPCR immediately.

The qPCR was executed on a Bio-Rad IQ5 with IQ SYBR, as described previously (38). Household genes Y37E3.8 and *rpl-6* were used as reference genes. αS and YFP were amplified for the detection of αS expression (**Supplementary Table S2**). After the Ct values were obtained, the relative abundance was calculated according to (38):

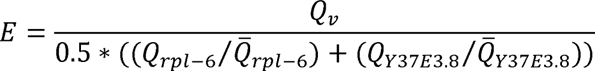

in this formula *E* represents relative abundance of target genes *v* (αS gene or the YFP gene), *Q* refers to the transformed abundance. The expression of target genes was normalized to housekeeping genes *rpl-6* and Y37E3.8 (38). These normalized expression values were used in further analyses.

#### Microarray expression normalization

Microarray data for the time-series, RIL and IL experiment were normalized with the Loess method for within-array normalization and the Quantile method for between-array normalization in R by Limma package (39,40). Afterwards, the log_2_ transformed intensities were used. Before analysis, we identified technical spots and spots that were associated with none or more than one transcript (based on a blast versus the WS258 *C. elegans* genome) (41). These 7,679 spots were censored in further analyses.

#### Principal component analysis (PCA)

The gene expression data was subjected to PCA aiming to explore the structure of the phenotypic variation in the αS-RILs as well as parental and reference strains. The PCA was conducted using log2 ration with the mean transformed gene expression data:

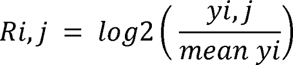

where R is the log_2_ relative expression of spot *i* (*i* = 1, 2, …, 45,220) in strain *j* (αS-RIL), and *y* is the intensity (not the log_2_-transformed intensity) of spot *i* in strain *j*. Subsequently, the *prcomp* function in “R” was used to calculate the principal components.

#### Heritability analysis

The log_2_-transformed intensities were used to calculate board-sense heritability (H^2^). H^2^ estimates were calculated as the fraction of phenotypic variance that can be explained by strain by estimating the technical variance from the repeats of the two parental and two genetic background strains. By estimating the genotypic variance and the remaining variance (*e.g.* measurement error) in the parental lines (as in (42)), heritability was calculated:

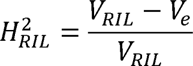

where *V_RIL_*is the variance within the RIL population and *V_e_* is the pooled variance of both parental and reference strains.

To establish whether the heritability was significant and not outlier driven, we applied a permutation approach (as in (43)). The trait values were randomized over the line designations and the heritability calculation were repeated. This was done 1000 times for each transcript to generate a by-chance distribution. The 50^th^ highest value was used as the FDR = 0.05 threshold.

#### Transgression analysis

Transgression was calculated by counting the number of lines with expression levels beyond three times the standard deviations of the mean from the parental strains (as in (44)). The lower boundary was established by the parental line with the lowest mean, and the upper boundary was established by the parental line with the highest mean. The standard deviation (σ) was calculated as the pooled standard deviation of the two parental lines (n =4 for both). We set the boundary for transgression at 3 * σ.

Significance of the transgression was calculated by permutation. The expression values were randomized over the line designations and the same test as above was conducted. This was repeated 1000 times for each transcript, so the obtained values could be used as the by-chance distribution. The 50^th^ highest value was used as the false discovery rate (FDR) = 0.05 threshold.

#### eQTL mapping

The eQTL mapping was done following the procedure as described previously(15,45). In short, we used a linear model to map gene expression QTL using a single marker model:

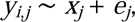

where *y* is the log_2_-normalized intensity as measured by microarray of spot *i* (*i* = 1, 2, …, 45220) of RIL *j*. This is explained by the genotype (either CB4856 or N2) on marker location *x* (*x* = 1, 2, …, 1927) of RIL *j*.

Permutations and calculations were performed to determine an empirical false discovery rate (45,46). We randomly permuted the log_2_-normalized intensities of each line per gene over the genotypes, which we used for mapping with the single marker model using the original genetic map. This was then repeated for ten randomized datasets as previously (45). The threshold was determined by a false discovery rate:

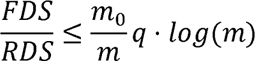

where FDS (false discoveries) is the permutation outcome and RDS (real discoveries) comes from the eQTL mapping at a specific significance level. The value of m_0_ and m were taken from the number of microarray spots (45220), representing true null hypotheses and hypotheses tested respectively. The FDR adjusted p-value (*i.e.* q-value) was set at 0.05 threshold.

Power analysis was done as described previously (15). In short, we simulated 10 QTL per marker location in normally distributed noise corresponding to various amounts of variance explained from 0.20, 0.25, …, 0.80. The data were subsequently mapped and analyzed at the previously determined FDR = 0.05 threshold.

#### eQTL analysis

The mapped eQTL were classified as either *cis*- or *trans-*regulatory. This was determined based on the location of the eQTL in correspondence with the regulated gene. If the gene was within the confidence interval of the QTL (based on a 1.5 drop in −log_10_(p) significance), the eQTL was classified as *cis*. In all other cases the eQTL was classified as *trans*.

The variance explained per microarray spot with an eQTL was calculated by ANOVA. For all spots, also those with multiple peaks, this analysis of the gene expression explained over the peak-marker was performed assuming a single-marker. For analytical purposes, we filtered the list of eQTL to a single representative spot per gene (the spot with the highest variance explained).

*Trans-*bands (an enrichment of *trans*-eQTL) were identified by counting a Poisson distribution of the mapped trans-eQTL (15,45,47). The number of *trans*-eQTL per 0.5 Mb block were counted, and then we calculated how many *trans*-eQTL were expected to be found. This we used to identify if an overrepresentation according to the Poisson distribution was present (p < 0.001). Thereby, *trans*-eQTL hotspots at specific markers were determined.

#### IL differential gene expression analysis

The IL experiment contained a detectable batch effect and therefore the expression was batch-corrected by fitting all the data per spot to a linear model and subtracting the batch effects. Thereafter, the expression effects attributable to introgression, presence of αS, and the interaction between the two were determined by:

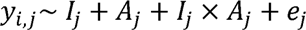

where *y* is the batch-corrected log_2_-normalized intensity as measured by microarray of spot *i* (*i* = 1, 2, …, 45220) of IL *j*. This is explained by the introgression *I* (either CB4856 or N2), the presence of the αS transgene *A* or the interaction between *I* and *A*. The model was fitted per set of strains (*e.g.* all the strains descending from the WN268 x NL5901 cross) and for each of the strains n = 3. Significance was determined by correction for multiple testing using the *p.adjust* function with the *fdr* method (q = 0.05).

#### IL – RIL comparative analysis

To determine the influence of the introgression, αS inclusion, and the interaction between the two, the effect sizes attributable to these terms were correlated with the eQTL effect sizes as measured in the RIL panel. For each of the IL sets (the groups of four ILs deriving from a specific cross) the genes with an eQTL mapping onto the CB4856 introgression locus were thus investigated. The Pearson correlation was calculated, as well as the significance of the correlation.

#### Enrichment Analysis

Gene enrichment analysis was done by a hypergeometric test on the genes, with Bonferroni-corrected *p<0.05*, and filtering for categories with at least three genes, with an overlap of at least two genes in the query. The following databased from www.wormbase.org WS258 were used: gene class annotations, GO-annotation, anatomy terms, phenotypes, RNAi phenotypes, developmental stage expression, and disease related genes (48). We also used the ModERN transcription factor binding sites from http://epic.gs.washington.edu/modERN/ as described previously (49).

#### Data analysis

Data was analyzed using RStudio (version 2021.09.0+351) with R (4.1.1, 64×). For data processing the tidyverse package was used (50) and plotting was done using the ggplot2 package (51). For some of the statistical analyses the ggpubr package was use (52). All data processing and analysis scripts were custom written and deposited at https://git.wur.nl/published_papers/huang_wang_2023_as_eqtl.

## Results

### Construction of the αS-RILs in N2 and CB4856 genetic backgrounds

In a previous study, we showed that transcriptome and fitness effects of αS introgression vary between genetic backgrounds (28). To be able to localize modifier loci, we crossed genetically distinct αS introgression lines (IL) SCH4856 and NL5901. These strains were chosen, because their parental genetic background (CB4856 and N2 respectively) are extensively characterized (14,53). Moreover, strain SCH4856 has more severe accumulation of αS that relies on protein accumulation rather than on differences in gene expression, meaning potential modifiers found in our approach are likely to affect protein aggregation which is more relevant to the human disease phenotype (54).

In total, 212 SCH4856 x NL5901 RILs were generated (hereafter referred to as αS-RILs). We randomly picked 90 of the 212 αS-RILs for genotyping by PCR (**Supplementary Table S1)** and subsequently 88 out of 90 for low-coverage whole-genome sequencing to generate a genetic map consisting of 1,927 markers (**Supplementary Table S3**; **Supplementary Figure S1**). A broad region in the center of chromosome IV and a small region on the right of chromosome X were completely N2-derived (**Supplementary Figure S1**). Given the larger size of the N2-derived fragment on chromosome IV region, this region was the most likely location of the αS insertion, although an insertion on chromosome X cannot be excluded. Practically these results implied that we will not be able to identify polymorphic loci residing in these fully N2-derived regions using (e)QTL mapping. However, our RIL population has sufficient coverage from both SCH4856 and NL5901 (**Supplementary Figure S1C**) on other locations and can be used to map genetic modifiers at other positions of the genome.

### Heritability and transgression analyses indicate a broad genetic basis for gene expression differences in αS-RILs

We set out to characterize the gene-expression variation in a time-series experiment on the two parental lines NL5901 and SCH4856 to determine the optimal time/age for taking samples to analyze gene expression differences caused by αS-genotype effect. We aimed to select a timepoint with few developmental gene expression differences between the parents as developmental effects may obscure effects caused by differences due to the αS phenotype. We found that at an age of 120 hours few developmental differences existed (**Supplementary Figure S2**). From the above-described NL5901×SCH4856 RIL panel a set of 70 sequenced RILs was randomly picked (**Figure 1A**). These were grown at 20 [for in total 120 hours, and for the last 72 hours kept on fresh FUdR plates. After 120 hours, the animals were collected, RNA was isolated, and gene expression was measured (**Figure 1B**).

**Figure 1:**
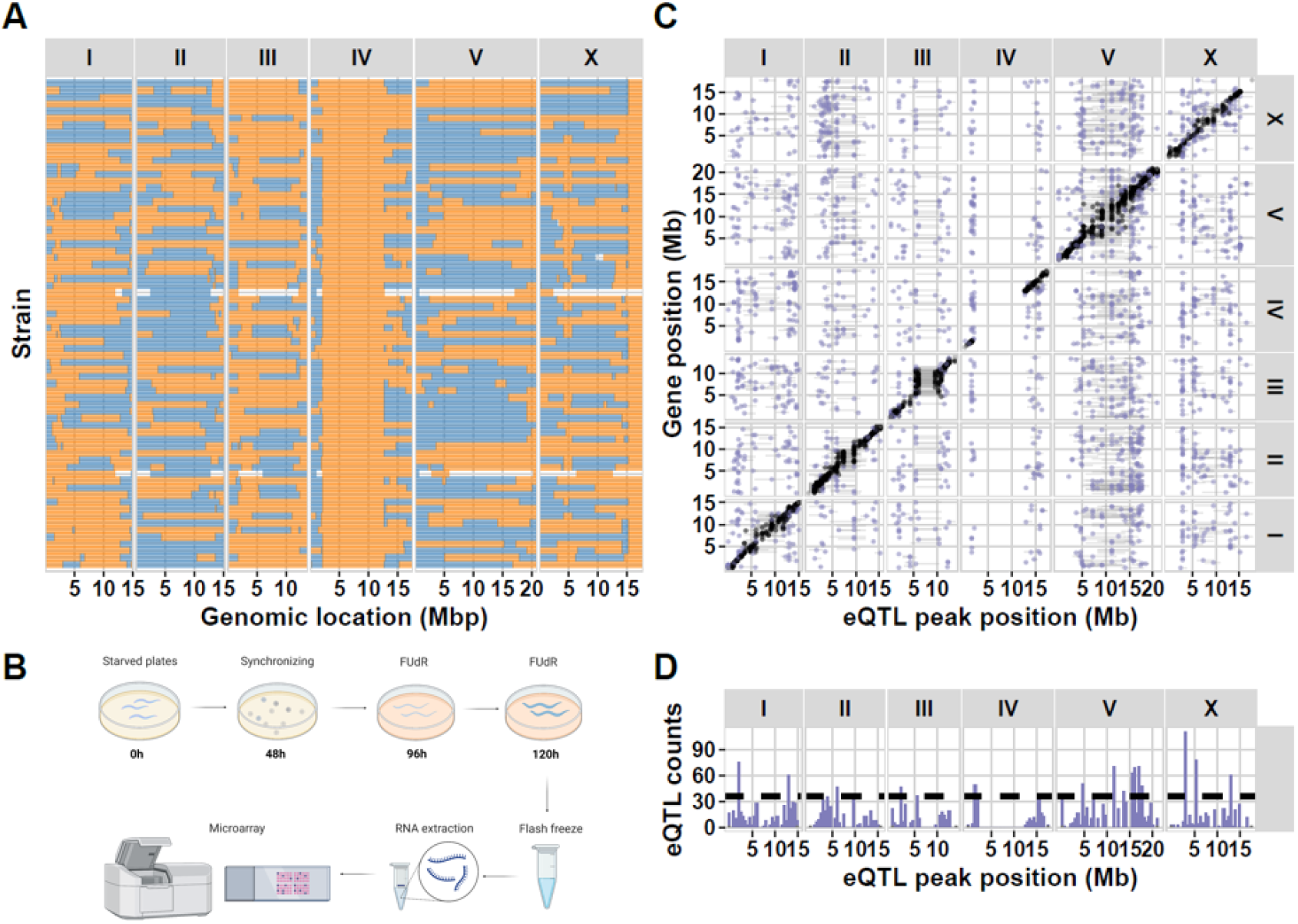
**(A**) The 70 NL5901×SCH4856 RILs used for eQTL mapping. (**B**) Experimental setup of the eQTL experiment from sample collection to gene expression measurements per microarray. (**C**) A plot of the *cis*- and *trans*-eQTL identified in the αS-RIL population. The x-axis shows the position of the eQTL and the y-axis the position of the gene that is regulated, both in million bases (Mb). The panels on the top- and right side indicate the chromosomes. Each dot represents the peak of an eQTL identified by mapping the expression data, the horizontal bars indicate the confidence interval of the eQTL. (**D**) A histogram indicating the density of *trans*-eQTL at each position on the chromosomes (per 0.5 Mb bin). The dashed horizontal line indicates the threshold for significant overrepresentation of *trans*-eQTL in a bin.

The genome-wide gene-expression was measured in these 70 αS-RILs and CB4856, N2, NL5901, and SCH4856 using microarrays. Initial principal component analysis (PCA) showed that the expression profiles of N2, NL5901, CB4856, and SCH4856 were clearly separated in accordance with their genetic background. Furthermore, the individual RILs were found in-between the SCH4856 and NL5901 strains (**Supplementary Figure S3**).

To determine the proportion of gene-expression variance caused by genetic variation in the population, we estimated the broad-sense heritability (*H^2^*) per microarray spot. We used the pooled variance of the four replicates of the four ancestral strains to estimate the technical variance (see Methods for details). The majority of significant *H^2^* values were in the range of 0.60–1.00 (**Supplementary Figure S4**). In total, 3,260 heritable genes were found (permutation, FDR = 0.05; **Supplementary Table S4**). This showed that the expression of many genes was heritable, hence there was a broad genetic basis underlying the variance in gene expression. To understand the genetic complexity underlying gene expression variance within the RIL population, we analyzed transgression by comparing gene-expression of the αS-RILs with both parental strains (NL5901 and SCH4856), identifying 1,468 genes showing transgression (permutation, FDR = 0.05) (**Supplementary Table S5**). This indicated that multiple loci from both parental strains were contributing to gene-expression variance. Interestingly, 621 genes were both heritable and transgressive (**Supplementary Figure S5**). These results together indicate that genetic variations interact with αS phenotypes.

### eQTL mapping in the αS-RIL population identified *trans*-eQTL hotspots on chromosome V

To understand the genetic basis for gene-expression variation, eQTL were determined in the αS-RIL panel based on genome-wide gene expression measurements. A power analysis showed that we could identify 84.2% of the eQTL explaining at least 35% of the gene expression variance. Using a single marker model, *cis-*eQTL (local acting, gene within the confidence interval of the QTL), *trans-*eQTL (distant acting), as well as eQTL hotspots could be identified. We identified 3,865 genes with an eQTL, including *trans-*eQTLs related to 2,406 genes and *cis-*eQTLs related to 1,827 genes (-log_10_(p)>4.3; permutation, FDR≤0.05) (**Supplementary Table S6**; **Fig 1C**). In the *trans*-eQTL set we identified 18 eQTL hotspots (Poisson distribution, p<0.0001; **Fig 1D**). These hotspots contained 1,167 genes, representing 48.5% of all *trans*-eQTL. Additionally, expression of αS was quantified by qPCR which showed that gene expression of this inserted gene was stable. Furthermore, no eQTL for αS expression was detected (**Supplementary Figure S5**). This implied that genome-wide gene expression differences were not due not due αS expression itself, but secondary effects, such as protein accumulation, which was observed before (Wang et al., 2019). In conclusion, we identified ample polymorphic genetic regulation of gene expression in the αS-RIL population.

Each chromosome was found to contain at least one eQTL hotspot (or *trans*-band) (**Figure 1D**; **Supplementary table S7**). These represent regions where the allelic variation among the RILs affects the expression of many genes at distant locations. In particular, we found that 8/18 *trans*-bands were located on chromosome V. Enrichment analysis showed that eQTL *trans*-bands were enriched for specific terms (**Supplementary table S7, Supplementary Figure S6**). For example, the chromosome V *trans*-bands were enriched for various gene ontology terms, mostly including terms related to development and transcription. We set out to independently verify the effects measured on chromosome V, as the *trans*-bands were only statistical genotype-phenotype associations.

### Chromosome V is associated with an up-regulation of *pmk-1*

To verify the effect of an αS introgression and natural genetic variation on chromosome V on gene expression, a set of ILs was constructed (**Supplementary table S8**). The region on chromosome V from 3.85 – 20.9 Mb, containing all eight chromosome V *trans*-bands, was covered by crossing three previously constructed IL_N2_ strains that contain a CB4856 fragment introgressed in an N2 background (WN268, WN269, and WN270) with NL5901 (Figure 2A) (14,29). From these crosses, four genotype sets per original IL_N2_ were obtained: i) harboring no αS and no CB4856 introgression, ii) harboring no αS but with CB4856 introgression, iii) harboring αS but no CB4856 introgression, and iv) harboring both αS and CB4856 introgression (Figure 2B). Thus, from the three crosses, in total 12 strains were constructed.

**Figure 2:**
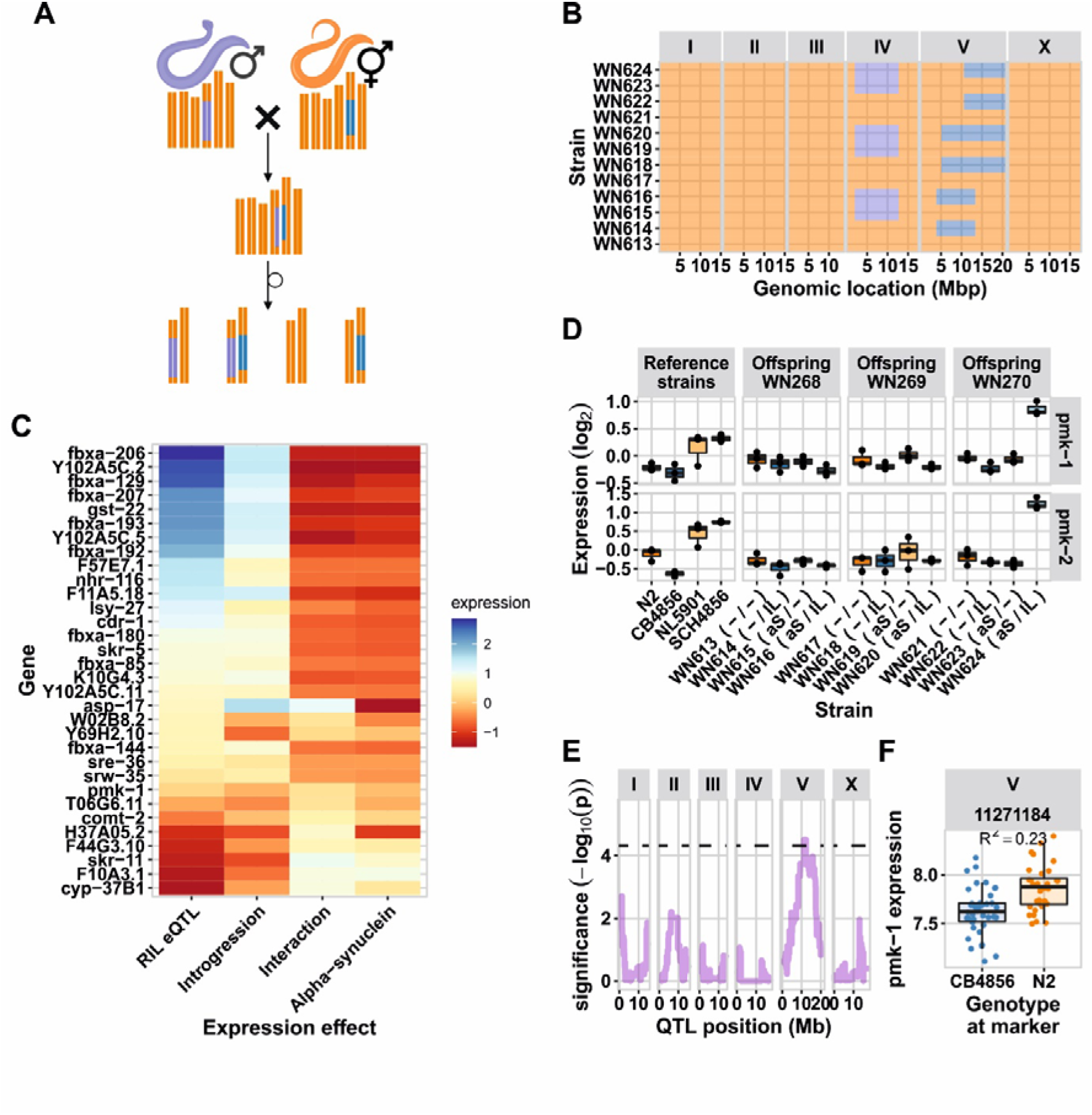
Introgression line experiments. (**A**) The crossing setup for the ILs, males of NL5901 were crossed with three different CB4856>N2 ILs. The F1 progeny were self-fertilized for 8 generations, ultimately leading to four types of strains. (**B**) The introgression line (IL) panel generated. Blue colors indicate the CB4856 genotype, orange the N2 genotype and purple the αS::YFP region on chromosome IV. (**C**) The relative expression of the genes with a significant interaction effect in the ILs with an eQTL on chromosome V. The RIL eQTL effect, and the expression in the ILs attributable to the introgression, the αS, and the interaction between the two are shown. Genes are ordered based on the RIL eQTL effect. (**D**) The expression of *pmk-1* and *pmk-2* in the IL experiment. The expression values shown are the log_2_ ratio with the mean. (**E**) The eQTL profile of *pmk-1* in the RIL experiment. The dashed horizontal line indicates the FDR = 0.05 threshold. (**F**) The genotype split-out at the eQTL peak, the presence of the CB4856 allele leads to lower expression.

To confirm the *trans*-bands and to determine whether the eQTL are the result of an interaction between the αS transgene and the CB4856-genotype, these αS-ILs were used for a gene-expression study similar to the αS-RIL experiment. Using correlation analysis, the eQTL effects as mapped in the RILs were correlated with the expression differences in the ILs attributable to CB4856 introgression, αS transgene, and the interaction between CB4856 introgression and αS transgene. The analysis showed that the introgression effect was dominant, explaining a significant amount of the measured *cis*- and *trans*-eQTL (*R* > 0.68, p < 1*10^-15^; **Supplementary Figure S7A**). We also found that a minor part of the expression variation in the *trans*-eQTL was affected by the interaction between the introgression and the αS transgene (|*R|* > 0.2, p < 1*10^-6^; **Supplementary Figure S7C**), whereas the αS inclusion itself was hardly associated with expression changes at the chromosome V locus (**Supplementary Figure S7B**). Altogether we conclude that the *trans*-bands on chromosome V are mainly the result from allelic variation, not from an interplay between αS and the CB4856 genotype.

Although the overall effect of introgression – αS interactions at the chromosome V loci was small, 74 genes with expression affected by the interaction were identified. Of these, 32 were mapped in the RILs as eQTL to the location of the introgression, most (29/32) as *cis*-eQTL (Figure 2C), and many of the genes fell into the *fbxa*-class. Upon case-by-case gene expression investigation of the genes with an interaction effect, we found that the *pmk-1* gene has been associated with late-onset PD through protection against cellular stress (Figure 2D) (55). We found that this gene also had an eQTL in the RIL panel (Figure 2E, Figure 2F). Both *pmk-1* and its close homologue *pmk-2* function in neuronal cells where they regulate neuronal responses and behavior (56). Although the observed expression effect is similar to the IL_WN268_ and IL_WN269_ sets (lower expression when the CB4856 allele is present), the IL_WN270_ set shows markedly high expression of *pmk-1* as well as *pmk-2* (Figure 2D). This indicates there are at least two regulatory loci underlying *pmk-1* and *pmk-2* expression, and these be highly upregulated due to a CB4856 locus on the right side of chromosome V.

## Discussion

### A novel RIL population to stud αS protein aggregation in the model nematode *C. elegans*

Identification of genetic modifiers of complex human diseases such as PD can provide crucial information about genetic mechanisms underlying disease and identify potential drug targets. Here, we constructed a new recombinant inbred line population in the genetic model organism *C. elegans* derived from strains NL5901 and SCH4856 with each strain carrying an αS human transgene (αS-RILs) to explore gene expression mechanisms underlying αS effects. Both strains were chosen because their parental genetic background (Bristol N2 and CB4856, respectively) have been well characterized (12). Furthermore, strain SCH4856 has a higher αS accumulation compared to NL5901 suggesting that potential background modifiers are likely to affect protein aggregation (54). To further characterize the underlying loci, a quantitative genetics approach using RILs and ILs was chosen (57).

The population of αS-RILs carries wild type alleles of Bristol N2 and CB4856 (58,59) and was constructed with the aim to avoid genetic imbalances and to achieve a large map size. Despite the efforts, significant allelic skews were found on most chromosomes. Most of these were in previously identified loci, namely: the *peel-1*/*zeel-1* locus on chromosome I (12,31) and the *npr-1* locus on chromosome X (12,60). Also, each αS-RIL has an N2 locus in the center of chromosome IV like the parental strains NL5901 and SCH4856. The genetic map showed that the αS gene was likely inserted at chromosome IV between 2.0 Mb up to 12.6 Mb but an insertion on chromosome X (15.1 Mb – end) cannot be excluded. In previous research, the αS transgene was estimated to be located on chromosome IV (4.2 - 13.2 Mb) (28). The differences in exact location, is mainly due to the less accurate estimation that could be provided in the previous study.

We were able to construct a population with a large map size. We used 88 sequenced αS-RILs to generate a genetic map consisting of 1,927 informative SNP markers which allows for studying quantitative trait variation for αS associated traits, complementing existing *C. elegans* models of αS. The map size was 120-210 cM per chromosome and in-between a RIL and a RIAIL population (61) and significantly larger than previously generated RIL panels used for studying specific mutations (15). Therefore, despite the allelic skews, the αS-RILs provide ample resolution to precisely map QTL.

### Ample quantitative genetic variation in gene expression in the αS-RIL panel

We previously found that presence of αS initiates gene expression that depend on the genetic background. In other words, depending on the genotype of the strain, gene expression responds differently to the insertion of αS (28). Therefore, we identified the effects of natural genetic variation and αS-inclusion on the whole-genome gene expression using an eQTL approach. Genetic modifiers in particular could change gene expression patterns genome-wide, thus having broad effects on how these different genotypes handle αS presence and potentially result in different disease outcomes (28). Therefore we built upon these findings to identify potential modifier loci.

The amount of heritable genetic variation and transgression was in line with previous observation in *C. elegans*. We find significant broad-sense heritability *H^2^* for 3,260 gene and for instance 2,910 genes were reported in a standard environment and 5,072 genes in a heat-stressed environment for a *C. elegans* N2 x CB4856 population of 54 RILs across three different temperature environments (Sterken et al. 2020). These results indicate that there is no major transcriptional effect due to the αS-inclusion at the time-point where expression was measured. However, our experiment was conducted in significantly older animals (120 hours old versus 48 hours old). Since gene expression regulation diminishes with age (43,46) we cannot exclude that the similar amount of genes with significant *H^2^* is affected by ageing.

Our eQTL mapping experiment identified a large number of *trans*-bands, many noticeably located on chromosome V. We found 1,827 genes with a *cis*-eQTL, 2,406 genes with a *trans-*eQTL, and 18 *trans-*bands. This is on the higher end compared with previous eQTL studies in *C. elegans* on similar microarray platforms, that report between 819 - 1,722 *cis*-eQTL and 816 - 1533 *trans-*eQTL (as deposited in WormQTL2.0) (15,41,45,47,62). This could indicate that the number of *trans*-eQTL increased due to genetic perturbations. However, the confounding effect of the genetic background (fixed on parts of chromosome IV and X), makes that from the αS-RIL panel alone, we could not be sure about potential modifier loci and needed an additional verification.

### Introgression lines for chromosome V point to a specific interaction between *pmk-1* and αS

To enable the detection of candidate modifier independent of the αS-RILs, we constructed ILs covering a large part of chromosome V. We found that *pmk-1* could be a modifier affecting αS in the αS-RILs. Previously, we discovered that *pmk-1* was affected by the developmental age of the nematodes with αS-carrying nematodes developing slower across genotypes (28). The protein coding gene *pmk-1* belongs to the MAP-kinase family and is homologous to human p38 MAPK (56). Although the exact function of (unaggregated) αS in humans is still unknown, it has been speculated that it may contribute to regulating cellular decision between autophagy and apoptosis with both protein degradation processes depend on p38 MAPK signaling (63). Since, experimental evidence in a *C. elegans* with 6-OHDA-induced PD has indicated that *pmk-1* is essential for effective treatment of PD effects using the flavonoid baicalin (64). Together, this suggest the p38 MAPK pathway forms an important target for PD treatments and our studies indicates that genotype-effects within this pathway may affect disease progression (28).

Our study is the first to explore genetic modifiers of αS in *C. elegans*. *C. elegans* has been extensively used as a model organism for studying protein accumulation, which is an important hallmark of NDG in humans. But most previous studies were limited to αS accumulation in a single genotype, i.e. the canonical strain Bristol N2, and did not allow to investigate natural genetic variants modifying the effect of αS. The current *C. elegans* αS-RILs provide a genetically diverse population which facilitates the investigation of polymorphic loci. Extending the quest of genetic variation studies promotes investigating the effect of αS in even more diverse wild type genotypes of *C. elegans* of which many are now available from CeNDR (Cook et al. 2017). Overall, these studies investigating genetic variation of NDG in model organisms will uncover genetic basis of disease, identifying key targets for human treatments.

## Author contributions

YAW, DV, YC and JAGR constructed the RILs. YH and JAGR constructed the ILs. YH, YAW, VT, and JAGR conducted microarray experiments. YH, YAW, JK, SCH, and MGS conceived the project. YH, YAW, LvS, and MGS analyzed the data. YH, YAW, LvS, JEK, SCH, and MGS wrote the paper; YAW began the project, YH finished it. All authors read and approved the final manuscript.

## Data availability

Data of the HiSeq sequencing for genotyping the RIL population was deposited at BioStudies under E-MTAB-12575. Transcriptomics data were deposited on BioStudies: the time series data (E-MTAB-11657), the RIL data (E-MTAB-11658), and the IL data (E-MTAB-12575). The created RIL and IL strains can be provided upon request.

## Acknowledgements

The authors want to thank Erik Andersen and Robyn Tanny for sequencing and analyzing the genotypes of RIL strains created in this study. Stefan van de Ruitenbeek is thanked for support with depositing the sequencing data.

